# Neuron types in the developing mouse brain are divided into partially matching epigenomic and transcriptomic classes

**DOI:** 10.1101/2022.12.29.522188

**Authors:** Sami Kilpinen, Heidi Heliölä, Kaia Achim

## Abstract

In recent single-cell -omics studies, both the differential activity of transcription factors regulating cell fate determination and differential genome activation have been tested for utility as descriptors of cell types. Naturally, genome accessibility and gene expression are interlinked. To understand the variability in genomic feature activation in the GABAergic neurons of different spatial origins, we have mapped accessible chromatin regions and mRNA expression in single cells derived from the developing mouse central nervous system (CNS). We first defined a reference set of open chromatin regions for scATAC-seq read quantitation across samples, allowing comparison of chromatin accessibility between brain regions and cell types directly. Second, we integrated the scATAC-seq and scRNA-seq data to form a unified resource of transcriptome and chromatin accessibility landscape for the cell types in di- and telencephalon, midbrain and anterior hindbrain of E14.5 mouse embryo. Importantly, we implemented resolution optimization at the clustering, and automatized the cell typing step. We show high level of concordance between the cell clustering based on the chromatin accessibility and the transcriptome in analyzed neuronal lineages, indicating that both genome and transcriptome features can be used for cell type definition. Hierarchical clustering by the similarity in accessible chromatin reveals that the genomic feature activation correlates with neurotransmitter phenotype, selector gene expression, cell differentiation stage and neuromere origins.

## Background

Clustering single cells by their transcriptome or chromatin accessibility allows to identify cell types and measure molecular distances between the cell types^1,2^. In the embryonic mammalian brain, the diversity of differentiating cell types is immense and largely unknown. Here, the topology of the cell type tree can provide a measure of developmental and evolutionary relatedness of cell types^3–5^: Specific patterns of genomic feature expression - or enhancer use - along the branches of a cell type tree can reveal genetic regulatory logic. For example, it has been shown that neurotransmitter phenotype can be associated with multiple TF combinations in *Drosophila* and *C. elegans*, and that the expression pattern of neural differentiation genes are better explained by a model where each gene can be regulated by several TFs^6,7^. The examples of such phenotypic convergence are still rare in vertebrates, however the divergence between lineage relatedness and final acquired cellular phenotype has been demonstrated in zebrafish lineage tracing studies^8,9^. Thus, similar molecular identity can be derived from molecularly and physically distinct lineage. The genetic elements underlying phenotypic convergence have this far only been studied in detail in C. elegans^10,11^.

Comparative analysis of genomic features is not always straightforward. While in RNA-sequencing features are defined by the consensus set of genes, this is not the case for the ATAC-sequencing today. In ATAC-sequencing, essentially reads could originate from any genomic area. In ATAC-seq data processing, features are formed *de novo* based on either i) fixed genomic windows or ii) locations of aggregated peaks of reads along the genome. The latter approach has clear benefit in increased resolution, but the results vary between samples. After successful feature definition, cell clustering and subsequent classification into cell types or transcriptomic classes is complicated by the variety of available clustering methods, as well as the lack of established cell type definitions and lack of comprehensive databases of marker genes for existing cell types. Additionally, correspondence between cell types defined based on transcriptome and chromatin accessibility profile should be considered. The within-cell-type diversity in both gene expression and chromatin accessibility can and probably does have different scales. It has been shown that clustering results using these different modalities do not perfectly match, however both yield valid classifications^3^. Finally, as in any clustering, cell type detection by single-cell data clustering is prone to bias due to under- or overclustering^12^. This risk can be mitigated by an iterative search for the resolution parameter that yields highest statistical confidence of clustering as well as by exploring biological interpretation for defined clusters.

Gamma-aminobutyric acid (GABA) is a small-molecule neurotransmitter and GABAergic neurons are the principal type of inhibitory neurons in the mouse brain. Consistent with the ubiquitous presence in the brain, GABAergic neuron precursors are found in all anterior-posterior divisions of the mouse neural tube, where they, interestingly, express and require distinct transcription factors (TFs) for acquiring GABAergic identity. Dlx1/2/5 genes function as GABAergic fate selectors in the telencephalon, Gata2, Gata3 and Tal2 in midbrain and Tal1 in ventral hindbrain^13^. The binding sites for above-mentioned fate selectors have been found in the regulatory regions of GABAergic neuron marker genes *Gad1* and *Gad2* (encoding glutamic acid decarboxylases 1 and 2 that convert glutamate to GABA in neurons)^14,15^. However, the gene regulatory elements underlying the flexibility to respond to various mutually exclusive developmental signals are not fully understood, neither the terminal differentiation genes possibly co-regulated with the canonical GABAergic neuron marker genes. To compare the gene regulatory landscape in different GABAergic neuron types arising from spatially distinct lineages during mouse embryonic development, we collected single cells from the embryonic day (E) 14.5, (E14.5) mouse telencephalon, diencephalon, midbrain and rhombomere 1. In the analysis workflow, we first define an ATAC-seq feature set by allowing separately collected single-cell populations to contribute into peak aggregation separately, then pool the peaks and finally use the resulting E14.5 common features to call accessibility of chromatin features per cell^16,17^. Next, we integrated scATAC-seq and scRNA-seq modalities using canonical correlation analysis (CCA)^17^. This allows using the dataset to study transcriptional and chromatin accessibility variation among cell types in an integrated fashion. In the integrated dataset, we studied the distribution and relatedness of cells in correlation with clustering, brain region neurotransmitter phenotype and selector gene expression. The expression of previously known regional markers in mouse brain as well as known GABAergic neuron fate selector genes aligned remarkably well with the scATAC-seq based clustering and could be correlated with novel regulatory DNA elements and RNA expression.

## Results

### Definition of open chromatin features in E14.5 mouse neural cells

To study transcriptional regulation of cell fate determination during neurogenesis, we first collected a diverse sample of neuroepithelial cells. Single cells were derived from the ventral telencephalon, diencephalon, midbrain and rhombomere 1 regions of a developing mouse embryo (Figure 1A). To map the open chromatin areas and the mRNA molecules expressed in the cells, we performed scATAC-seq and scRNA-seq on stage- and region matched samples (Figure 1B, Methods).

**Figure 1.**
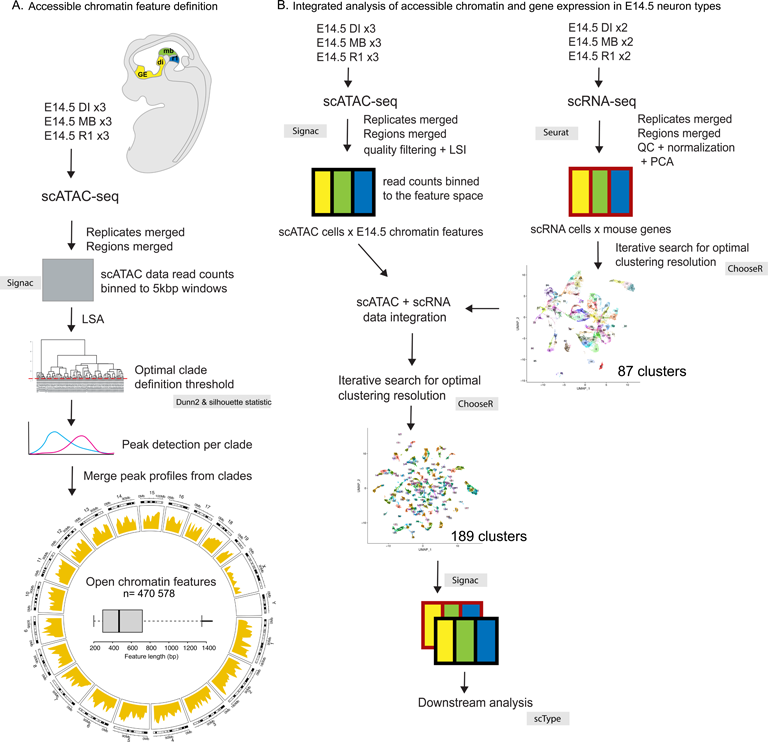
Graphical overview of the presented workflows. A. Accessible chromatin feature definition. The analysed brain regions are listed and shown in colour on the schematic of mouse embryo. Samples were collected from the telencephalon (Tel) and diencephalon (Di), midbrain (MB) and rhombomere 1 (R1) regions of E14.5 mouse embryos and used as indicated in each workflow. The process for defining the ATAC feature space across studied brain regions is described, showing the methods and steps of the workflow. See Methods for details. The circosplot shows the density of features (yellow) across the mouse karyotype. Boxplot shows the distribution of features by feature length in base pairs (bp). Outliers (>Q3 + 1.5*IQR) were excluded from the boxplot. B. Schematic of the scRNA-seq and scATAC-seq data processing and integration workflow. Samples (number of replicates indicated) were subjected to single-cell mRNA-seq (scRNA-seq) or single-cell ATAC-seq (scATAC-seq). Each replicate consists of ca 5000 cells (scRNA-seq) or nuclei (scATAC-seq) from a single embryo. The sequencing reads from scRNA-seq (red outlines) or scATAC-seq (black outlines) are first processed in parallel, then combined into an integrated dataset. The scATAC-seq data is scored along the previously defined E14.5 ATAC features. scRNA-seq data is processed and collapsed into reads per gene level. After clustering resolution optimisation using ChooseR, the scRNA-seq data is integrated with scATAC-seq using Seurat/Signac, followed by the optimisation of the clustering based on ATAC features (ChooseR). Cell type classification (scType) is done after the integration of scATAC-seq and scRNA-seq data. Tel, telencephalon (yellow); Di, DI, diencephalon (yellow); MB, midbrain (green); R1, rhombomere 1 (blue). LSA, Latent Semantic Analysis; QC, Quality Control; PCA, Principal Component Analysis; scATAC, scATAC-seq, single-cell ATAC-sequencing; scRNA, scRNA-seq, single-cell mRNA-sequencing.

The sampled brain regions are enriched in various subtypes of GABAergic neurons (see Methods). In addition to GABAergic neurons, the sampled brain regions contain neuroepithelial progenitors, glutamatergic and cholinergic neurons, and monoaminergic neuron types. To compare the gene expression and regulation across all the cell types, we decided to pool the samples. Analysis of the set of pooled cells originating from different scATAC-seq experiments required features applicable to all the samples. We addressed this by merging the reads from the separate scATAC-seq experiments, and then defining features stepwise: we first used the reads from cells that formed high-level clades separately^16^. To determine optimal cutting level for clade detection we used Dunn2 and Silhouette statistics^18^ (Supplementary Figure 1A-B, Figure 1A). After clade definition, we performed separate peak detection in each of the clades. Clade level peak definition was followed by peak merging (Methods) resulting in a final set of n= 470 578 features (Figure 1A, *E14.5 ATAC features*).

We further characterized the defined features in terms of feature length, density, and localization to genomic context. Features were found to be mostly under 1000 bp long and covered entire mouse genome (mm10) with numerous hotspots throughout the studied genome (excluding X- and Y-chromosomes) (Figure 1A). Altogether, the defined features cover approximately 8.7% of the mouse genome sequence (236 594 772 bp out of total of 2 730 855 475 bp). Nearly 30% of the features are located within +/− 10kb around TSS, which altogether is a relatively narrow part of the genome. Approximately 80% of the features locate within +/-100kb around TSS, containing both genes and intergenic regions (Figure 2A). Average distribution of the feature locations over gene models from TSS to TTS (transcript termination site) clearly peaks at the TSS but also shows that regions downstream of TTS have slightly more features (Figure 2B). This highlights that while TSS regions are, as expected, the most dynamically regulated areas of chromatin, significant stretches of intergenic chromatin are accessible in at least some cellular clade. We annotated the features by the genomic region and compared with randomized positions of equal-length fragments (Figure 2C), confirming highest enrichment of features in proximal promoter regions (<1 kb of TSS) and introns (Supplementary Table 1, Figure 2C). Features are significantly less often found in intergenic regions (Figure 2C, Supplementary Table 1). In total, 45% of the E14.5 ATAC features are found in intergenic regions while 55% are located within gene structures.

**Figure 2.**
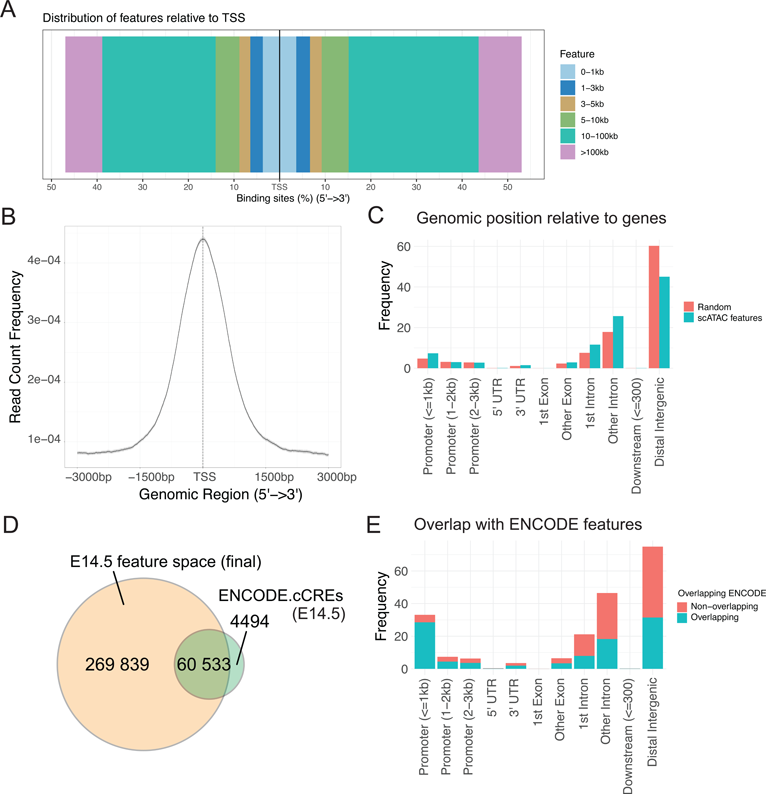
Characterisation of the E14.5 ATAC features. A. Distribution of the E14.5 ATAC features relative to the TSS (transcription start sites) of genes in mouse genome. X-axis shows cumulative percentage of features locating upstream or downstream of the TSS. Colors indicate distance intervals from TSS in kb. B. Count frequency of feature peak locations along the average profile of gene from TSS to TTS (5’-> 3’ direction). C. Bar chart of all E14.5 ATAC features (blue) annotated for the genomic position relative to genes. Random position (red) is the average proportion for randomly positioned features, calculated using 1000 random shuffles of peak locations, per annotation category. D. Overlap between the features in the E14.5 mouse forebrain, midbrain and hindbrain according to ENCODE (ENCODE cCREs) and the expression-filtered E14.5 ATAC features used in the downstream analysis. E. Genomic regions of the expression-filtered E14.5 ATAC features and their overlap with the ENCODE features (blue). The features not overlapping with the ENCODE features are shown in red.

To assess how each sample can be represented using the E14.5 ATAC features, we compared the open chromatin features defined in each brain region separately with the common E14.5 ATAC features. 94-99% of the features present in the separately analyzed samples fully overlap a feature from the common E14.5 ATAC features (Supplementary Figure 1C for E14.5 DI). In all brain regions, 0.4 - 3% features remained region specific (Supplementary Figure 1E). The common E14.5 ATAC features also contain features not found in any brain region alone (Supplementary Figure 1D-E). Likely, combining reads from the samples enhances the signal from weakly detected common features. We did not detect significant differences in the contribution of reads from individual samples to E14.5 ATAC features (Supplementary Table 2). In conclusion, we cannot see meaningful bias between samples in their contribution to defined E14.5 ATAC features.

Features used in the downstream analysis were further filtered based on detected accessibility in at least 1% of the cells and in maximum of 97.5% of the cells, resulting in 330 384 chromatin features. This set of expressed E14.5 ATAC features was compared to the functional DNA elements available from ENCODE ^19^ for E14.5 forebrain, midbrain and hindbrain (See methods for ENCODE sample ID-s). 93.1% of the ENCODE elements are found in our feature set (60533/65027 elements). However, our E14.5 features contain 269839 additional features, making the ENCODE elements a minority (15.48%) of the defined features. Aside from the noise inherent to single-cell data, our feature set may be enriched for rare features that are expressed transiently and not to a significant extent in the cell type level. The features overlapping with ENCODE are more often found in promoters (Figure 2E), for example 28.51% of features overlapping with ENCODE are ≤1kb from promoters versus 4.58% of non-overlapping features.

### Cell type identification and comparison between brain compartments

We next used the defined chromatin features to analyse the scATAC-seq data from E14.5 DI, MB and R1 together (Figure 1B). After filtering the scATAC-seq cells by having at least 2.5% of features and maximum of 97.5% of features accessible, the E14.5 scATAC-seq dataset comprised 330 384 chromatin features in 28 505 cells (Figure 1B, left side). Subsequently, we integrated the E14.5 scATAC-seq data with E14.5 scRNA-seq data (Figure 1B, right side). First, the cells from the scRNA-seq samples from E14.5 DI, MB and R1 were combined, expression was scored across mouse genes and cells were clustered (Methods). The resolution parameter for cluster detection was estimated by iterating cluster detection over range of resolution values and selecting the highest statistically supported resolution using *chooseR* ^12^. The clustering resulted in 87 clusters at the optimal resolution (Supplementary Figure 2, Methods). We did not observe significant batch effect in the scRNA-seq samples (Supplementary Figure 2). The scATAC-seq data and scRNA-seq data integration was performed as described by Hao *et al*. ^20^ and Stuart *et al.* ^21^. The final integrated dataset contains 20 606 cells.

After the modality integration, we proceeded to cluster the E14.5 cells based on their scATAC-seq profile. The optimal resolution parameter for cluster detection was res=16, as defined using *chooseR* ^12^ (Supplementary Figure 3, Supplementary Table 3). The scATAC-seq dataset was divided into 189 cell clusters (Figure 3A). The cells from the replicate embryos were distributed evenly across the clusters (Supplementary Figure 4C, G). We did not observe clustering by the genotype of the collection date of cells (Supplementary Figure 4D, H), indicating the lack of significant batch effect. The “Di” samples from batch 2 (06.10.20) contain diencephalic and telencephalic cells and therefore may group by brain region separately from the diencephalon samples from batch 1 (06.08.20, embryo 1). This sample group is referred to as Di+Tel in further text and Figures. The cells from each brain region mostly formed coherent clusters (Figure 3B, Supplementary Figure 4B, F), with some mixed clusters that may contain migratory cells present near midbrain-hindbrain boundary, or cells near regional borders that may be found in differently labelled samples.

**Figure 3.**
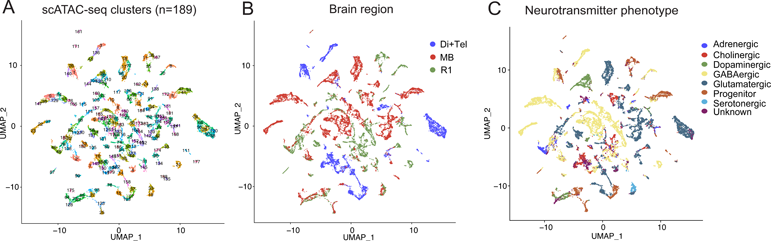
Clustering of the E14.5 Di/Tel, MB and R1 cells based on the scATAC-seq feature expression. A. UMAP projection of the clusters of E14.5 cells. Clustering is based on the scATAC-seq feature accessibility. B. UMAP of the clusters showing the brain region of origin of each cell. C. UMAP of the clusters showing the identified neurotransmitter phenotype of the cells.

To characterize the cells in each cluster further, we assigned a neurotransmitter phenotype or progenitor cell identity to each cell, implementing *ScType*^22^. We first defined a set of positive and negative marker genes for the cell types present in the diencephalon, mid- and hindbrain at E14.5 (Supplementary Table 4). Those include GABAergic and glutamatergic neurons that are derived from ventrolateral and lateral neural tube, dopaminergic, serotonergic and cholinergic neurons lineages derived from floor plate and basal plate, and adrenergic neurons of the R1. The definition of glial cells (astrocytes and oligodendrocytes) was not included in *ScType* and thus the possible glial cells present in the sample were likely placed in *unknown* category (1631/20606 cells, 7.9%, Supplementary Table 4). Overall, we observed high level of concordance between cluster identity and neurotransmitter phenotype (NT-type, Figure 3C), with NT-type purity value ^23^ being 0.87 (Methods). To analyze the enrichment of known neuronal marker genes or housekeeping genes among the differentially expressed genes in the clusters, we calculated Gini index ^24^. Mean Gini index was 0.81 for the neuronal marker genes and 0.12 for the housekeeping genes, indicating that housekeeping gene expression is relatively uniform, whereas expression of neuronal marker genes is variable across the clusters. Consistent with the NT-type results, the markers for the cell types present in the diencephalon, mid- and hindbrain at E14.5 were differentially expressed in clusters (Figure 4). The final number of scATAC-seq cell clusters (n=189) matches well with the expectation to find the 6 neuron classes as well as the neuronal progenitors, while each neuron type can be present in several stages of neurogenesis and maturation in E14.5 mouse brain, which is further divided into dorso-ventral (7 domains) and anterior-posterior (4-5 domains) molecularly distinct domains and may give rise to temporal sublineages^25–27^. Importantly, the categorization by neurotransmitter identity reveals broad classes of neurons, while distinguishing the whole range of molecular identities would require an improved set of markers. ScType also do not directly support dual neurotransmitter identities, which therefore may have been placed in the “unknown” category or receive a poor NT-type score. This may be the case for GABA and acetylcholine dual neurotransmitter cell clusters in the diencephalon scRNA-seq samples (Supplementary Figure 5A, Supplementary Figure 6B-C, see below). As we had sampled several cell lineages per neurotransmitter identity in each brain region (Supplementary Table 5), we defined the clusters further, finding combinatorial marker gene sets using *CombiROC* ^28^ (Methods, Supplementary Table 6). In all brain regions, the several cell clusters classified as the same neurotransmitter phenotype, were characterized by a unique marker gene combination (Supplementary Table 6). To validate the specificity of the marker gene combinations, we analysed the sampled cells for the co-expression of all genes in the found cluster marker lists (152/189). Considering each marker combination, the averaged expression of all genes in the marker list was calculated per cell. In the calculations of average expression, the cells were filtered so that only cells showing expression value > 0.98 quantile of the given gene over all cells were considered. This was done to reduce background and visualise cells expressing the marker genes at high level. The expression of each cluster marker combination was visualised and can be found at https://github.com/ComputationalNeurogenetics/NeuronalFeatureSpace/.

**Figure 4.**
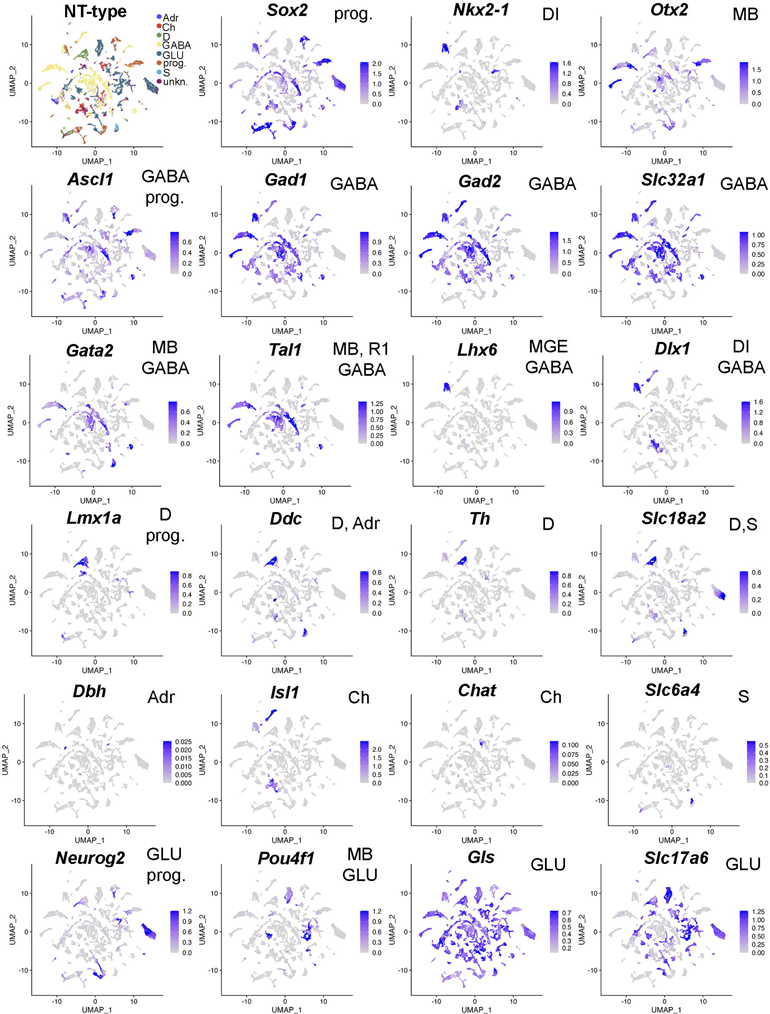
Neurotransmitter phenotypes (NT-type) and the related marker gene expression in the E14.5 mouse brain. Expression of known neuronal marker genes and selected transcription factors plotted over the UMAP projection of cell clusters. Intensity of blue color indicates the relative level of gene expression. The shown marker genes used in the NT-typing (see Supplementary Table 3). The respective cell type is indicated at each plot. prog, progenitor cells; GABA, GABAergic neurons; D, dopaminergic neurons; Ch, cholinergic neurons; Adr, adrenergic neurons; S, serotonergic neurons; GLU, glutamatergic neurons; MB, midbrain; R1, rhombomere 1; DI, diencephalon.

We next wanted to explore the relationships between the cell types. For that aim, we calculated average chromatin accessibility profile for each cell cluster and subjected those to hierarchical clustering (Figure 5, Methods). Interestingly, hierarchical clustering by chromatin accessibility revealed some segregation of clusters by brain region of origin as well as by neurotransmitter type (Figure 5). However, the brain region and NT-type do not correlate fully, and there seems to be no clear preference to group by either characteristic. Surprisingly, several cluster groups comprised of a single cell type segregate very early: a branch of Di+Tel glutamatergic neurons (Di Glut; Group 9) appeared at 3^th^ branching event (Figure 5C, and Supplementary Figure 5A-C), and midbrain glutamatergic neurons (MB Glut, Group 2) at 4^th^ branching event (Figure 5C, and Supplementary Figure 5). A midbrain GABAergic neuron group appears at branch 6 (MB GABA, Group 5, Supplementary Figure 5). Diencephalic ACh+GABA neuron group at branch 7 (Supplementary Figure 5A, Group 5). A dopaminergic neuron group (DA) of mixed sample origin (Di and MB) appears at 8th branching event (Figure 5; Supplementary Figure 5, Group 6). The DA neurons locate in diencephalon and midbrain in the mature brain, while the DA neuron progenitor domain is in ventral midbrain, and thus the genetic regulation of dopaminergic fate is similar. The first branching event in cluster tree is clearly associated with the separation of progenitor cells from the other, more differentiated groups (Figure 5, Supplementary Figure 5A, prog), however some progenitors are retained in both branches. Some branches of mixed cell types of single brain region origin can be seen (Figure 5), however at least in early branching the mixed clusters tend to remain mixed both in cell types and regions. It would be interesting to study how the chromatin accessibility signature is predictive of cell type or region. Most interestingly, similar cell types appeared at multiple locations in the hierarchical tree. Such branches may represent cell types with different chromatin accessibility configuration that yet can acquire a similar cellular phenotype. We explored this phenomenon further by analysing the clusters comprising GABAergic neurons (see below).

**Figure 5.**
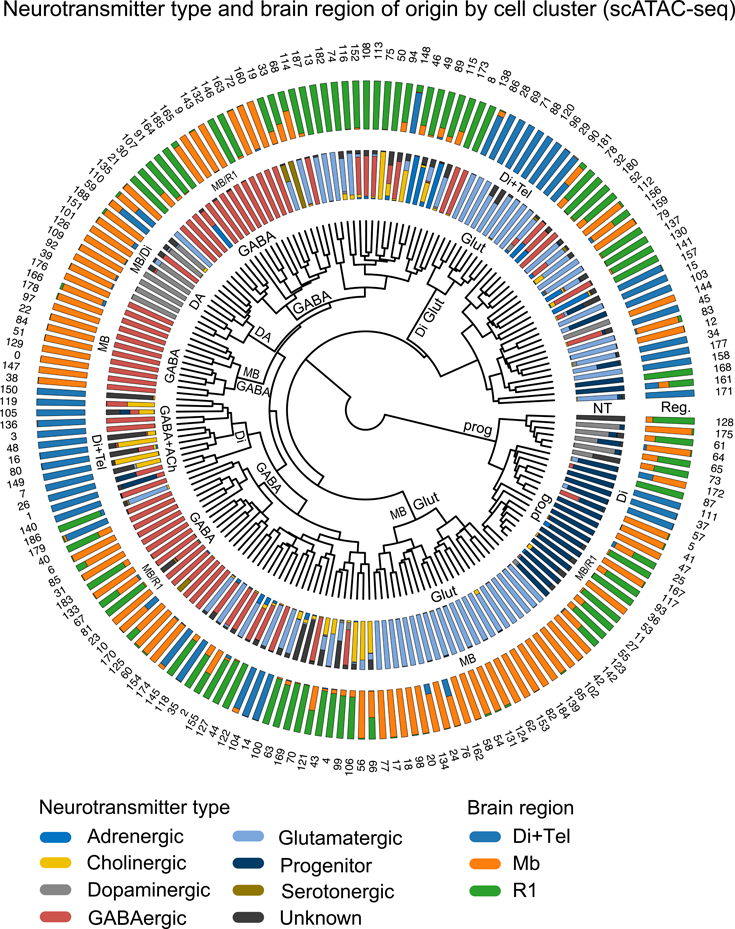
Analysis of the clustering by cell attributes. A. Circos plot, showing starting from inside: the dendrogram of E14.5 cell clusters (n=189) based on E14.5 ATAC feature accessibility, the neurotransmitter type per cell (NT), the brain region of origin (Reg.), and cluster numbers. Stacked barplots show the proportion of the cells derived from different brain regions or the proportion of cells of different neurotransmitter identity per cluster. Labelling in text indicates the branching events where a cluster of cells of single cell type or region of origin emerges. NT, neurotransmitter type; Reg, brain region identity. Brain regions: Di +Tel, diencephalon and telencephalon; MB, midbrain; R1, rhombomere 1. GLUT, glutamatergic neurons; GABA, GABAergic neurons; DA, dopaminergic neurons; prog, progenitor cells.

### Similar cluster identities are found using transcriptome and chromatin accessibility as distance measure

The accessible chromatin areas affect gene expression and thus largely define boundaries for RNA expression in the cells. To explore the similarity between the cell clusters formed based on accessible chromatin regions or based on mRNA expression, we calculated the cluster-to-cluster matches between the scATAC-seq and scRNA-seq cell clusters (Methods, Supplementary Figure 6D). We then compared the distribution of neurotransmitter phenotypes between the clusters (Supplementary Figure 6C-D, Supplementary Table 5). In the matching clusters, the cell types and the brain region of origin were mostly similar (Supplementary Figure 6D). There are some important differences between the modality specific clustering results^2^. As our scATAC-seq data contains more clusters, the cells clustering by scRNA-seq often matched cells from several scATAC-seq clusters (Supplementary Figure 6D). It is likely that the resolution reached here does not allow definition of all the cell types or cellular stages. For example, we did not identify dominantly adrenergic or cholinergic clusters in the scRNA-seq (Supplementary Table 5). This might be improved by sampling more cells.

### Clustering of GABAergic neuron precursors in E14.5 mouse brain correlates with the selector gene expression and differentiation stage

We next focused our analysis on the GABAergic neuron subtypes. The predominantly GABAergic clusters were defined as clusters containing >75% of cells with NT-type=GABAergic and cluster mean silhouette reliability score > 0.5. We selected these clusters (n=52) and studied their relatedness in terms of chromatin accessibility profiles in isolation. Hierarchical clustering of the GABAergic neurons resulted in a complex tree structure in terms of brain region of origin (Figure 6A). We then asked how the clustering correlates with the known GABAergic neuron marker gene expression. As expected, the GABAergic neuron marker genes *Gad1* and *Gad2* were expressed in all the identified GABAergic neuron clusters (Figure 6A). During development, GABAergic neuron precursors express proneural transcription factor *Ascl1* and a combination of selector transcription factors *Dlx1/2, Gata2/3* and *Tal1/2* that are required for the acquisition of GABAergic phenotype. We observed clear differential expression of *Ascl1*, *Tal1, Tal2, Gata2, Gata3, Dlx1* and *Dlx2* correlating with the clustering (Figure 6A). Consistent with previous studies, the clusters originating from diencephalon and telencephalon (Di+Tel) expressed *Dlx1* and *Dlx2*, with the exception of cluster 10 that expresses *Gata3* and thus represents the derivatives of the rostral prosomere 2 (P2R) or prosomere 1 (P1) region of the diencephalon^29^. The cells originating in midbrain and rhombomere 1 expressed *Tal1*, *Tal2*, *Gata2* and *Gata3* in various combinations. The selector genes for the dorsal and dorsolateral (D/DL) R1 GABAergic lineages are unknown^30^. Interestingly, while the GABAergic clusters of Di, MB and the D/DL R1 appear to group, the branches also share similarity in differentiation stage. We find two mixed groups of differentiating neurons (expressing *MAPT* and *Tubb3*), early GABAergic neuron precursors (expressing *Tal2, Tal1, Ascl1*) of MB and two branches of *Tal1/2, Gata2/3* positive, intermediate GABAergic neuron precursors of mixed origin in MB or R1. Finally, clusters from all three brain regions are found in the *MAPT+* differentiated neuron branch. In conclusion, the grouping of the GABAergic neuron clusters shows a loose correlation with the region of origin, differentiation stage and selector gene expression. While early precursors group according to the positional identity, at the transition to committed precursor stage, the cells of distinct lineages may display a transient variation in genome accessibility, which leads to similar priming of GABAergic differentiation genes. This phenomenon has earlier been characterized as phenotypic convergence ^31^.

**Figure 6.**
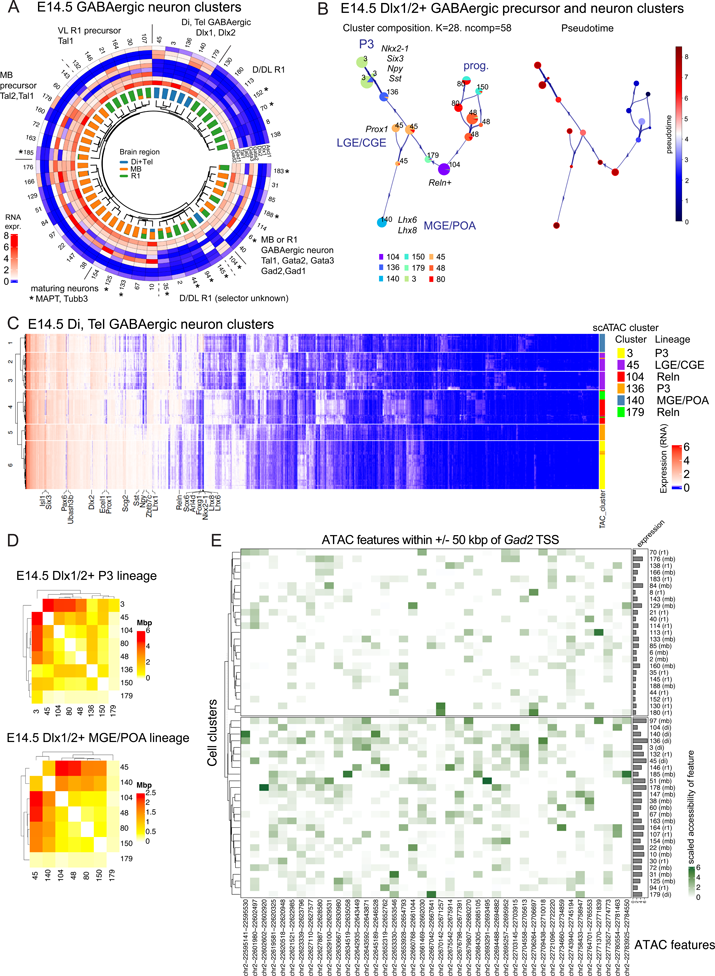
Differential chromatin accessibility associated with the GABAergic neuron lineages in different brain regions. A. Circos plot showing the hierarchical clustering dendrogram of dominantly (>75%) GABAergic clusters (n=52) in E14.5 mouse brain, barplots of proportion of cells by the brain region of origin, and the heatmap of average expression of the proneural gene *Ascl1*, GABAergic fate selector genes *Dlx1, Dlx2, Gata2, Gata3, Tal2* and *Tal1* and the GABAergic neuron marker genes *Gad1* and *Gad2*. The clustering by chromatin accessibility loosely correlates with the brain region and the selector gene expression. D/DL R1, dorsal/dorsolateral R1; VL, ventrolateral. Asterisk indicates the cell clusters where *MAPT* and *Tubb3* are found among the top 25 of differentially expressed genes. B. Chromatin accessibility based pseudotime trajectory and placement of the *Dlx1* and *Dlx2* expressing GABAergic cell clusters and progenitor cell clusters 80, 150 and 48. Cluster subdivision performed internally by VIA are shown on the left side, with colored pie charts indicating composition of each subcluster, where the numbers indicate original cluster number having majority in the corresponding subcluster. Lines with arrows indicate lineage relations and direction between nodes. The image on the right side shows the same lineage structure, with each subcluster colored according to estimated pseudotime. The red-framed circles indicate the lineage end points. The size of the circles is proportional to the number of cells in the cluster. The lineages representing the diencephalic prosomere 3 (P3), medial ganglionic eminence/ preoptic area (MGE/POA) and lateral and caudal ganglionic eminence of telencephalon (LGE/CGE) GABAergic neurons are indicated, along with the relevant marker gene expression. C. Heatmap of the inferred RNA expression in *Dlx1/2* expressing GABAergic neuron clusters. Hierarchical clustering is based on the expression of top 1400 most variable genes. Both axes have been clustered with Euclidean distance and ward.D2 linkage. scATAC-seq cluster of origin and the lineage placement of cluster is shown on the right for each cell. D. Heatmap of the total length of the DA chromatin in *Dlx1/2* expressing sublineages. The cell clusters were clustered by the similarity in the differentially accessible (DA) DNA features. E. Heatmap of chromatin accessibility of features within +/-50 kpb around *Gad2* gene (columns) across GABAergic clusters (rows). Dominant brain region of origin is shown in the parenthesis after cluster number. Barplots on the right side show average expression of *Gad2* in the cluster.

To further study the open chromatin features associated with the cell types and with the transition from an early precursor to late precursor and differentiating neuron stage, we applied pseudotime analysis using VIA ^32^. We focused on the *Dlx1*, *Dlx2* expressing GABAergic clusters (n=6; Figure 6A, Di, Tel GABAergic). The *Dlx1/2*+ clusters 45, 3, 136, 140 and 179 form one branch of the DA feature-based hierarchical clustering tree, with one *Dlx1/2*+ cluster (104) grouped with differentiated R1 neurons (Figure 6A). These 6 clusters with addition of 3 *Dlx1/2+* progenitor clusters 45, 48 and 80 (Methods) were selected for pseudotime analysis (Figure 6B). In pseudotime trajectory, three distinct lineages could be identified: the medial ganglionic eminence and preoptic area GABAergic neurons (cluster 140, Figure 6B, MGE/POA), the lateral and caudal ganglionic eminences group (cluster 45, LGE/CGE) and the diencephalic prosomere 3 lineage formed from clusters 3 and 136 (Figure 6B, P3). The MGE/POA clusters are positive for *Lhx6, Lhx8, Nkx2-1, Foxg1* expression, the LGE/CGE for *Prox1* and the P3 clusters by *Nkx2-1, Six3*, *Lhx1, Sst, Npy* RNA expression (Figure 6B-C). As the *Dlx1/2* expressing clusters did not fully group in chromatin accessibility based hierarchical clustering, we asked how much variation these clusters show in mRNA expression. Hierarchical clustering based on 1400 (5% of mouse genes) highly variable genes in the *Dlx1/2+* clusters show that cells similar in chromatin accessibility mostly cluster based on their transcriptome as well (Figure 6C). The cells forming lineages in pseudotime analysis also clustered based on mRNA expression (Figure 6B-C, *Reln+* clusters 104 and 179). In conclusion, we observe both differences of genome accessibility in transcriptionally similar cells as well the mRNA expression differences in the scATAC-seq clusters. The clusters that appear similar on the level of genome-wide chromatin accessibility can be further characterized for placement in distinct cellular lineages and for transcriptomic profiles.

The differentially accessible (DA) chromatin between the *Dlx1/2+* GABAergic clusters is ranging from 250-8000 kbps (Figure 6D). To ask how the DA chromatin contributes to the gene regulation, we applied Gene Ontology (GO) term enrichment analysis using genes nearest to the differentially accessible DA features in the neighbouring cluster pairs in pseudotime (clusters 45 and 3 and clusters 45 and 140, Figure 6B, Supplementary Table 7). In the GO biological process category, the DA feature-associated genes were greatly enriched in genes involved in neuronal development (*regulation of neuron migration, dendrite morphogenesis, axon guidance, homophilic cell adhesion via plasma membrane adhesion molecule)* or TF function *(positive regulation of transcription by RNA polymerase II*). Enriched cellular component terms (*neuronal cell body, postsynaptic membrane, presynaptic membrane, dendrite, dendritic spine, axon, growth cone, integral component of postsynaptic density membrane, neuron projection*) and molecular function terms (*cell-cell adhesion mediator activity, glutamate receptor activity, transmitter-gated ion channel activity involved in regulation of postsynaptic membrane potential*) were also associated with neuronal development and function or with TF function (*RNA polymerase II cis-regulatory region sequence-specific DNA binding, DNA-binding transcription activator activity, RNA polymerase II-specific*) (Supplementary Table 7). In summary, the differential chromatin accessibility in developing forebrain GABAergic neurons clusters is associated with the processes of neuronal differentiation.

Finally, we explored the differential use of regulatory elements near the GABAergic neuron differentiation genes. Presynaptic GABAergic neurotransmission requires maintained expression of *Gad1, Gad2* and *Slc32a1*. To understand how these genes are regulated in overall differing genome accessibility context in MB/R1 and Di/Tel, we mapped the feature accessibility near the *Gad2, Gad1* and *Slc32a1* genes in the GABAergic neuron clusters, showing that the feature accessibility increases in parallel with the target gene mRNA expression and observing high variability in feature accessibility 50 kbp up- and downstream of *Gad1, Gad2* and *Slc32a1* (Figure 6E, Supplementary Figure 7). Surprisingly, although the Di/Tel and MB/R1 tend to cluster in hierarchical clustering (at least considering *Gad2* and *Slc32a1* loci), we observe high level of variability in feature accessibility in all GABAergic neuron subtypes, both between and within the brain region. It would be interesting to study the common and differentially used genomic features for the selector TF binding.

## Discussion

### Cell type and state

Which modality, chromatin accessibility or the mRNA expression, is a better predictor of cell identity? Due to both technical and biological variability among single cells, cell type should not be defined as a transcriptionally and epigenetically fixed entity, but the signature should allow certain level of variation. Our cross-modality cluster comparisons confirm that we can detect different transcriptional and epigenetic states within cell types. Compared to the number of genes, the higher number of open chromatin features allows to reach higher clustering resolution. High resolution was achieved here by combining two essential steps: first, forming cell clades before feature definition allows to detect and consider rare and cell-type specific features, and second, using the highest clustering resolution achievable for given features and number of cells. Indeed, the number of clusters detected in the scATAC-seq experiment is nearly double compared to the number of clusters detected in the scRNA-seq data at optimal resolution. This has earlier been interpreted as segregation of different cell states ^2^. Alternatively, the one-to-many matching effect between scRNA-seq and scATAC-seq clusters could be due to the use of alternative gene regulatory programs. In this case, rather than cell state, the apparent ‘resolution’ reflects parallel regulatory or co-regulatory events, leading to similar mRNA expression profile. Careful analysis of marker gene combinations is required to address this question.

### The advantages and limitations of the study

As open chromatin features are stage- and tissue specific to a considerable extent, our study provides a pertinent method to combine data from different experiments. Nevertheless, additional experiments may be required to understand the regulation of cell differentiation. The chromatin features defined in this study cover 8.7% of mouse genome and can be used for the analysis of regulatory landscape and its dynamics in the E14.5 mouse CNS. However, using predefined areas for scoring chromatin openness has several limitations. First, dynamic use of chromatin outside of predefined features is not considered. Second, we lack information about the function of the feature in genome regulation (steric or regulative interactions). For the gene regulatory features, defining the target genes is not trivial. Feature-to-target gene links are especially challenging to define for the distant intergenic features (>100 Mb from genes). Such enhancers can associate with the target gene promoter by chromatin looping and their target genes can be either up-or downstream of the feature. Third, in the integrated scRNA-seq+scATAC-seq data, we characterized the cell clusters using the RNA expression inferred from a stage- and region-matched dataset. Nevertheless, rather than the inferred RNA expression level, true multimodal -omics data would allow to better assess the match between the RNA expression-based clustering vs clustering by open chromatin features. On the other hand, the mRNA expression response from chromatin accessibility change might require some distance in time ^3,33^.

Finally, mRNA expression and open chromatin features describe only a fraction of the cell. Post-transcriptional aspects such as the alternative splicing of RNA could not be addressed in this study. scRNA-seq also lacks information of the regulatory RNA molecules and non-polyadenylated RNA, such as long non-coding RNA. Providing a true snapshot of the genetic regulation network in a cell would also require in-parallel information of protein expression in the cells. Currently available methods limit the proteomics and protein-protein interaction studies as well as studies of post-translational regulation of protein expression and activity in a comparable cell type resolution.

### Regulative genomic features in mouse CNS development

Regardless of the limitations, our results and data comprising cells of numerous types and developmental stages will be an essential addition to the studies of neurogenesis and early fate specification of neurons. Developmental stages of active neurogenesis are under-represented in the current brain cell atlases.

In this study, we have demonstrated that using chromatin features for clustering single cells, a high resolution where ‘cell types’ are separated by hundreds to thousands of base pairs difference in accessible chromatin can be reached. Within GABAergic neurons, the similarity in chromatin accessibility correlates with cell subtypes and differentiation stage. Both GABAergic (as well as glutamatergic) neuron subtypes show remarkably distinct accessible chromatin profile in different brain regions. It is yet unclear how the observed difference in accessible chromatin is regulative to cell fate decisions. Cell fate is regulated by inter- and intracellular signalling that is, to a large extent, mediated by transcription factors. Transcription factor binding profile in differentially accessible chromatin features could therefore be indicative of developmental history, as well as predict the course of differentiation. The transcription factor and RNA polymerase binding at promoters and enhancers is accompanied by the modification of histones. Histone modification profile could thus be utilized as supporting evidence of gene regulation. Importantly, we have here integrated the chromatin feature accessibility and the mRNA expression data in single cells. mRNA expression adds multiple layers of value, first, it provides readout of gene transcription, and second, information about the molecular function of the cell or cell type.

Our comparison of the differentially expressed genes and the genes near DA features showed that the achieved resolution allows extraction of regulatory events associated with cell state, progression along differentiation stages, or even specific molecular functions. Furthermore, the enhancer usage could be associated with the target gene expression in GABAergic neurons.

### Conclusions

We have demonstrated an approach to study divergent gene expression regulation events in cell types within and across regional boundaries in mouse brain. The study can easily be extended to various questions in developmental biology, such as the transcriptional regulation of gene expression during neurogenesis and differentiation. We demonstrate how specific points of the “cell type tree” can be analyzed in detail for the appearance of cell types. In the future, it would be interesting to dissect the fractions of chromatin specifically associated with distinct but interlinked developmental events.

## Methods

### Single-cell RNA-sequencing

Single cells were isolated from *Nkx2-2^Cre/+^; R26R^TdTomato/+^* mouse embryos^34^. *R26R^TdTomato^* allele was obtained from Jackson laboratories, catalog number 007909). Embryonic day 0.5 (E0.5) was defined as noon of the day of the vaginal plug. On the selected embryonic day, the pregnant mouse was sacrificed by CO_2_ exposure followed by cervical dislocation, and embryos were dissected.

For the scRNA-seq, brains were isolated from E14.5 mouse embryos, and the ventral and ventrolateral forebrain, midbrain and R1 pieces were separated. The dissected tissue is relatively enriched in neuromeres producing GABAergic neurons. Lateral and medial ganglionic eminences and the preoptic area were extracted from forebrain, and the prosomere 3 and ventral parts of prosomere 1 and prosomere 2 of diencephalon. M2-M7 domains were collected from the midbrain. From rhombomere 1, the ventral and ventrolateral tissue was extracted, and the rhombic lip excluded. The area boundaries at zona limitans intrathalmica, midbrain-hindbrain boundary, alar-basal plate boundary and pallial-subpallial area border were identified by anatomical landmarks. Cells were dissociated using the Papain cell dissociation kit (Worthington, LK003150). Chromium 3’ single-cell RNA-seq kit (10xGenomics, 1000146) was used for cell capture and cDNA synthesis, with the cell yield set at 5000 cells per sample. Per each brain region, single cell collection was performed from two individual embryos. cDNA libraries were sequenced on Illumina NovaSeq 6000 system using read lengths: 28bp (Read 1), 8bp (i7 Index), 0 bp (i5 Index) and 89bp (Read 2). The reads were quality filtered and mapped against the Mus musculus genome *mm10*, using CellRanger v3.0.1 pipeline for scRNA-seq. The exact cell number, read yields and read quality for each sample are found in (https://github.com/ComputationalNeurogenetics/NeuronalFeatureSpace/seq_quality_reports).

### Processing of scRNA-seq data

scRNA-seq data processing per brain region is described in detail in R code in (https://github.com/ComputationalNeurogenetics/NeuronalFeatureSpace) with main steps outlined below.

scRNA-seq data was read into *Seurat* objects from 10x *filtered_feature_bc* matrix by using Ensembl gene IDs as features. Replicates per each brain region, or all replicates in the later analysis steps were merged as *Seurat* objects. After merging, quality control (QC) was performed by calculating *mt-percentage*. Cells were filtered requiring *nFeature_RNA* > 2500 and *nFeature_RNA* < 6000 and *percent.mt* < 15. Data was then log-normalized, top 3000 most variable features were detected, data was scaled and *percent.mt*, *nFeature_RNA*, *nCount_RNA* were set to be regressed out.

Data was then run through PCA using variable features, and *Seurat RunUMAP* with PCA dimensions 1-37 was used to generate Uniform Manifold Approximation and Projection (UMAP) for further analysis. *Seurat* functions of *FindNeighbors*(dims=1:37) and *FindClusters*(resolution=5) were applied for neighborhood and community detection-based clustering. Clustering resolution was optimized with *chooseR* ^12^. Neurotransmitter type was estimated by using *ScType* ^22^ as described below.

### Single-cell ATAC-seq

Embryonic brains were isolated from E14.5 wildtype (n=2) and Pax7*^Cre/+^; R26R^TdTomato/+^* mouse embryos (n=1). *Pax7^Cre^* allele (catalog number 010530) was obtained from The Jackson Laboratory. From each embryo, the ventral forebrain and diencephalic prosomere 3 (DI), ventral midbrain (MB) and ventral R1 pieces (R1) were separated and processed as individual samples. Nuclei were isolated using the recommended protocol for scATAC-seq (Assay for Transposase Accessible Chromatin, 10xGenomics protocol CG000212). At least 2 replicates were collected per each brain region and stage. Chromium Single Cell Next GEM ATAC Solution (10X Genomics, 1000162) was used for the capture of nuclei and the accessible chromatin capture. In chip loading, the nuclei yield was set at 5000 nuclei per sample. Chromium Single Cell Next GEM ATAC Reagent chemistry v1.1 was used for the DNA library preparation. Sample libraries were sequenced on Illumina NovaSeq 6000 system using read lengths: 50bp (Read 1N), 8bp (i7 Index), 16bp (i5 Index) and 50bp (Read 2N). The reads were quality filtered and mapped against the Mus musculus genome *mm10* using Cell Ranger ATAC v1.2.0 pipeline.

### Definition of feature space of scATAC-seq data

In order to define feature space that could be used in the processing of scATAC-seq samples from different brain regions, we adapted the approach described by Cusanovich *et al.* ^16^.

scATAC-seq data from the replicates of the same brain region were processed through pipeline described in (https://github.com/ComputationalNeurogenetics/NeuronalFeatureSpace) by using *Seurat* and *Signac* packages in R ^20,21^, obtaining read count tables per cell per 5kbps bin over the entire mm10 genome (excluding X- and Y-chromosomes). These were merged into one *Seurat* object and additional QC was performed based on nucleosome signal, TSS enrichment, blacklisted regions and reads in peaks fractions, as well as setting min and max limits for number of cells having count per feature and number of features having count per cell.

We then binarized the data and used Latent Semantic Analysis (LSA) ^16,21^ to reduce the dimensionality. The first singular component was omitted as it significantly correlated with read depth (Supplementary Figure 3C). Remaining 49 components were used to construct hierarchical tree of cellular clades with using cosine distance and ward.D2 as linkage methods. Next, we computed Dunn2 and Silhouette statistics ^35^ at various cut heights of the cellular clade tree (see Supplementary Figure 1A) by using R package *fpc*. Silhouette statistics measures how well each cell fits into the assigned cluster as comprared to other clusters. Dunn2 statistics measures separation or distance between clusters. Based on these statistics we selected cutting height h=6 having highest for both statistics, which resulted in in 74 cellular clades, with 65-1847 cells per clade (Supplementary Figure 1B). First, the reads from each clade were processed separately, using *Macs2* for the peak detection, thus each clade had equal possibility to provide its own peaks to merged collection ^16^. Subsequently, peaks from all clades were merged to form common E14 ATAC features^16^.

### Feature comparisons and characterization

In the comparisons of sample-specific features in VENN diagrams, the features were considered overlapping when they shared an overlap of at least of 100 bp. Features of the joined feature space were further characterized by using *ChIPseeker* ^36^ and visualized with *circlize* ^37^ R packages. Comparison of genomic annotation for peak locations for joined feature space vs random space was done with 1000 random iterations of peak locations and averaging proportions of hits to various genomic annotation categories. Fisher’s Exact test was used to test significance of observed vs expected proportions of annotation categories.

### ENCODE elements comparison

Comparisons of our features to known mouse genome elements were calculated with *findOverlapsOfPeaks* function from *ChIPpeakAnno* R package ^40^, by using parameters (connectedPeaks = “keepAll”, minoverlap = 1). The previously known elements were obtained from ENCODE database ^19^ from SCREEN UI V3 using closest matches in terms of brain region and embryonic stage: for E14.5 Di we used E14.5 mouse forebrain data, for E14.5MB we used E14.5 mouse midbrain and for E14R1 we used E14.5 mouse hindbrain data (ENCODE sample IDs ENCFF274NPS, ENCFF475FUT, ENCFF081VBW, MB_ENCFF539CBL, ENCFF829ZIT, ENCFF104QAC, ENCFF237YNH, ENCFF349REI, ENCFF279EVI).

### Processing of scATAC-seq data with E14.5 ATAC features

E14.5 ATAC features were used to process the scATAC-seq data as previously described, but instead of fixed 5 kbps windows, the features of the joined feature space were used. Data was subjected to QC, binarization and LSA, steps described in more detail in (https://github.com/ComputationalNeurogenetics/NeuronalFeatureSpace) largely following steps described by Stuart et al. ^17^. scATAC-seq data was integrated with scRNA-seq using an approach described by Hao *et al.*^20^.

UMAP projection based on scATAC-seq data was done with *Seurat* by using SVD components 2-59 for cells with reliable score from scRNA-seq integration. First component was omitted due to the association to read depth (Supplementary Figure 3C). Neighborhood and community detection were made with Seurat functions of *FindNeighbors*(dims=2:59) and *FindClusters*(resolution=16). Clustering resolution was optimized with *ChooseR* ^12^ as described below.

### Clustering resolution optimization

We used *ChooseR* ^12^ package as wrapper to test clustering resolution parameter values between 1-20 for clustering of both scATAC-seq and scRNA-seq data before visualizing with UMAP. We modified *ChooseR* package code to accommodate scATAC-seq data by adding an option to skip first SVD component in scATAC-seq data. The optimal resolution parameter was chosen based on silhouette values (Supplementary Table 6, Supplementary Figure 2).

### Identification of neurotransmitter type

Neurotransmitter types for each cell were identified by using *ScType* ^22^ from which we returned cell level typing, instead of more typical cluster-level cell type label, and we added an ‘unknown’ label based on a cell not having score for any defined cell type higher than mean(scores) + (1.3*sd(scores)). *ScType* was applied on integrated RNA data in scATAC-seq object and separately to RNA data in scRNA-seq object. The neurotransmitter phenotype of the cell types was identified based on the expression of positive and negative marker genes describing each neurotransmitter type (Supplementary Table 4). In addition to the different neurotransmitter types, we created ‘progenitor’ label using artificial composite genes called *g2m* and *s*, which describe the cells in G2/M or S-phase of the cell cycle, respectively. We adjusted a *gene_set_prepare* function in *ScType* to accept our added artificial composite genes (code in https://github.com/ComputationalNeurogenetics/NeuronalFeatureSpace). Otherwise, the code was run with default parameters. Marker genes for G2/M and S-phase were collected from *Seurat* cell cycle gene collection. As some of these genes are not entirely cell cycle specific in the context of developing neurons in mouse, we applied additional filtering by excluding genes showing expression in postmitotic neurons (Supplementary Table 4).

### Chromatin accessibility-based similarity of cell clusters and transcriptome correspondence

Similarity of scATAC-seq based cell clusters was further analyzed by hierarchical clustering. The average accessibility profile across all features per cluster was calculated and the clusters subjected to hierarchical clustering using cosine distance and ward.D2 as linkage method. The 189 identified scATAC-seq clusters were visualized with *circlize* with additional information of brain region of origin and neurotransmitter types visualized in stacked barplots.

In analysis focusing to GABAergic cell clusters, we selected cell clusters where >0.75 proportion of cells received NT-type=‘GABAergic’ label from ScType. These clusters were subjected to separate hierarchical clustering (cosine distance and ward.D2 linkage) based on their average accessibility profile across all features. The tree and selected cell attributes were visualized with *circlize*.

### Analysis of cell attributes distribution on the hierarchical cluster tree

Relative enrichment (R) of attribute categories (NT-type, brain region) was calculated at each branch up to ten branching points. R=((number of cells with the category/number of cells in total in that branch)/(total number of cells with the category/total number of cells)). R=1 indicates no enrichment of the particular category among the cells of the branch in question, whereas R value less or more than one indicate lower or higher relative proportion of the cells with that label, respectively.

### Pseudotime analysis

We used generalized trajectory inference method VIA ^32^ to examine trajectories and pseudotime among prominently GABAergic cell clusters based on chromatin accessibility. Additional three *Dlx1, Dlx2+ NT-type = ‘progenitor’* clusters (80, 48, 150) were added to the analysis to provide progenitor cells and root the lineages. Details and parameters of VIA analysis are available from (https://github.com/ComputationalNeurogenetics/NeuronalFeatureSpace). Data subset was exported from R and subjected to VIA analysis in Python (Jupyter) based workflow where VIA was run with *pyVIA* library. Progenitor clusters were set as initial roots.

### Unique marker gene combinations for cell clusters

CombiROC ^28^ was applied find unique combinations of expressed genes per each scATAC-seq based cluster. The thresholds for each cluster were decided based on visual consideration of plots provided by CombiROC. Thresholds and code are available from https://github.com/ComputationalNeurogenetics/NeuronalFeatureSpace NT purity and Gini index

To assess the biological validity of clusters, we calculated purity value ^23^ across clusters, as well as Gini index^24^ of expression of housekeeping genes and genes expressed in neurons. Both indexes were calculated based on imputed RNA. For both indices, the maximum value of 1 would indicate perfect segregation of NT-types/marker gene expression among the clusters ^39^

### scATAC-seq vs scRNA-seq cell cluster composition

In order to assess similarities in scATAC-seq and scRNA-seq cell cluster composition, for every cell, the cluster ID vectors of scATAC-seq clusters (n=189) and label transferred scRNA-seq clusters (n=87) were registered. The resulting contingency table (Supplementary Table 8) was visualized in heatmap with *ComplexHeatmap* package ^41^. Ordering of the rows and columns in the heatmap was calculated with R package *seriation* ^39^ by using BEA_TSP as method.

### Characterisation of differentially accessible chromatin

Pairwise differentially accessible chromatin features between all pairs of clusters were identified with *Seurat FindMarkers* function applying *min.pct=0.05*, logistic regression and latent variable set to *peak_region_fragments*. This was run in wrapper across all cluster combinations and differential accessibility (DA) was defined when avg_log2FC > 0.75 or avg_log2FC < −0.75 and p_val_adj < 0.05.

The closest gene for each DA feature was identified with *Signac ClosestFeature* with threshold of avg_log2FC > 0.7 for DA feature. Enrichment for the genes in Gene Ontology (GO) biological process categories was calculated with go_*enrich* function from R package *GOfuncR* ^38^.

### Accessibility of features around a selected gene

Cluster averaged and column scaled accessibility values were drawn for all features within selected window (+-50kpb from TSS) around the gene in question. Heatmap has been clustered with Euclidean distance and ward.D2. linkage methods. Row-wise partitioning has been found with k-means clustering and by searching for optimal k with maximizing silhouette score.

## Declarations

### Ethics approval and consent to participate

This research involves materials collected from laboratory mice, following the ethical research guidelines given by EU in *Directive 2010/63/EU on the protection of animals used for scientific purposes*. All animal experiments were approved by the Laboratory Animal Center, University of Helsinki, and the National Animal Experiment Board in Finland. The study was conducted and reported in accordance with the ARRIVE 2.0 guidelines (https://arriveguidelines.org).

## Consent for publication

Not applicable

## Availability of data and materials

The E14.5 rhombomere 1, midbrain and diencephalon scRNA-seq and scATAC-seq reads generated in this study are available in SRA under BioProject PRJNA929317. All original code will be deposited in GitHub at https://github.com/ComputationalNeurogenetics/NeuronalFeatureSpace Any additional information required to reanalyze the data reported in this paper is available from the corresponding author upon request.

## Competing interests

The authors declare that they have no competing interests.

## Funding

This research was supported by the Academy of Finland, decision numbers 319906 and 345737.

## Authors’ contributions

SK and KA conceptualized the study, planned the experiments, wrote the manuscript and compiled the figures. SK contributed to the formal analysis, implementation of methods, manuscript writing and editing. HH implemented the ScType for the classification of cells based on neurotransmitter phenotype. HH applied CombiROC for combinatorial expression marker search and participated in formal analysis. KA contributed to single cell RNA-seq and single cell ATAC-seq experiments, interpretation of results, manuscript writing and editing, supervising the project and acquisition of funding.

## Acknowledgements

We thank Institute of Molecular Medicine Finland Single-cell Genomics Core Facility, Jenni Lahtela and Bishwa Ghimire for the advice and assistance in single-cell sample processing and data analysis. We thank Juha Partanen and Janne Ravantti for reading and commenting the manuscript.

---

**Supplementary Figure 1.**
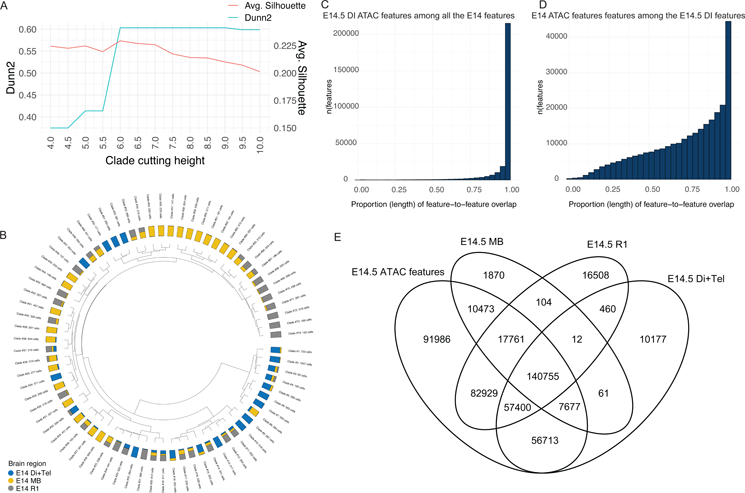
Distribution of cells in clades and the comparison of the features defined from individual scATAC-seq samples with the joined feature space. A. Dunn2 statistic (green) and cluster average Silhouette statistic (red) for each configuration of clusters acquired by cutting cellular tree from heights 4 to 10, with 0.5 step. The method seeks for maximum of both values, and minimal cutting height where distinct cell populations are separated. B. Dendrogram of cellular clades after applying the cutting height 6. Stacked barplot in second track shows the proportion of cells derived from each brain region. Outermost track shows the clade numbers and the number of cells per clade. C. Histogram of accessible features in the E14.5 DI sample that overlap with the features in E14.5 feature space, stratified by the proportion of overlap. D. Histogram of E14.5 features that overlap with the features defined using only E14.5 DI (E14.5 DI features), stratified by the proportion of overlap. E. Venn diagram of feature correspondences between separately defined and the E14.5 ATAC features.

**Supplementary Figure 2.**
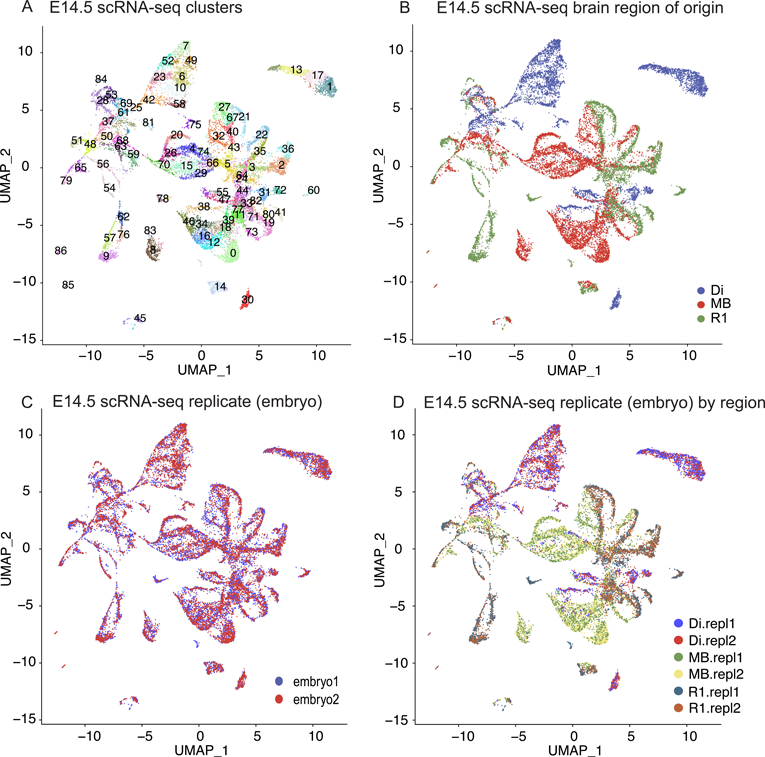
Batch effect and quality control analysis of E14.5 scRNA-seq samples. A. UMAP projection of clusters (n=87) of E14.5 scRNA-seq data. B. UMAP projection of clusters, cells colored based on brain region of origin. C. UMAP of scRNA-seq clusters, cells colored based on the replicate (embryo). C. UMAP of scRNA-seq clusters, cells colored based on replicate/embryo and region of origin.

**Supplementary Figure 3.**
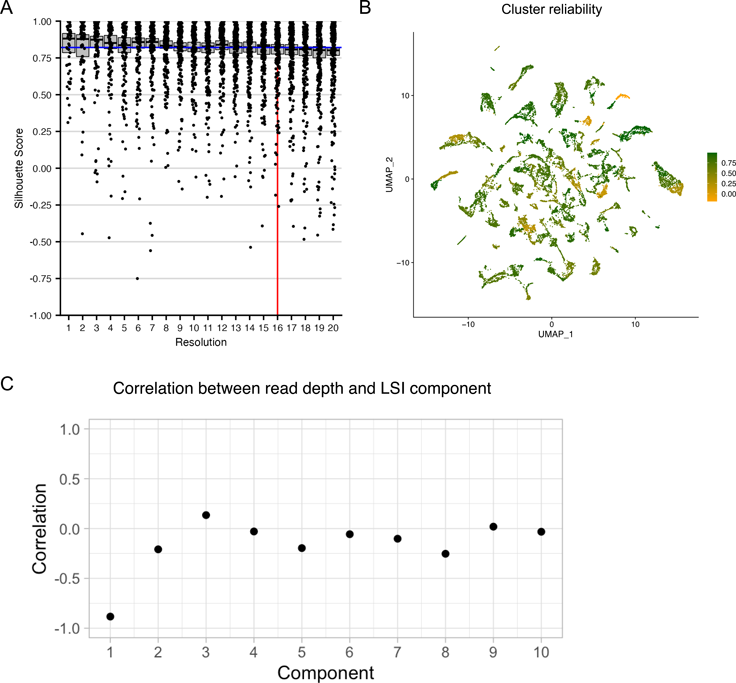
Clustering resolution, cluster reliability and the similarity of labels after the scRNA and scATAC based clustering. A. Silhouette statistic scores per clustering resolution calculated using *chooseR*. Optimal resolution is indicated with red line. Each dot represents at cluster. See the Supplementary Table 6 for the number of clusters at each resolution from res 1-20 and other details. B. Silhouette scores per cluster overlayed on the UMAP projection of clusters. Higher values (max 1) indicate clusters with high reliability. In further analysis, the representativeness of cell types should be carefully considered for clusters with the average silhouette score of <0.5. C. Correlation between read depth and LSI component. High (r<-0.5 or r>0.5) positive or negative correlation indicates components to be excluded from further analysis.

**Supplementary Figure 4.**
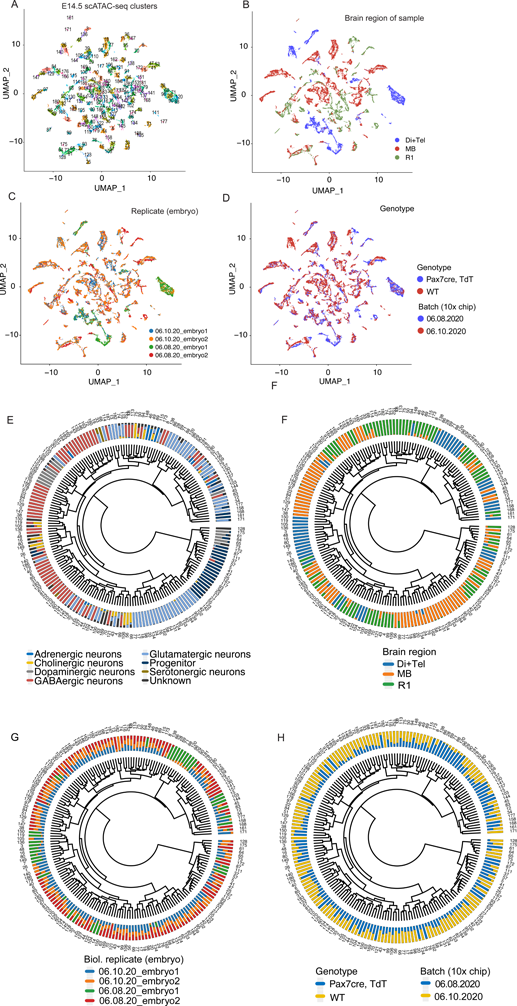
Batch effect and quality control analysis of scATAC-seq samples. A. E14.5 scATAC-seq clusters (n=189) on UMAP. B. Cells on the UMAP colored based on brain region of origin. C. Cells on the UMAP colored based on replicate (embryo). D. Cells on the UMAP colored based on genotype of mouse. E. Circosplot of all E14.5 scATAC-seq clusters (same as Fig. 5), with the proportion of cells in each cluster by the NT-type shown as additional track. F. Circosplot of all E14.5 scATAC-seq clusters, with brain region of origin proportions shown. G. Circosplot of all E14.5 scATAC-seq clusters, with replicate (embryo) proportions shown. H. Circosplot of all E14.5 scATAC-seq clusters, with genotype proportions shown.

**Supplementary Figure 5.**
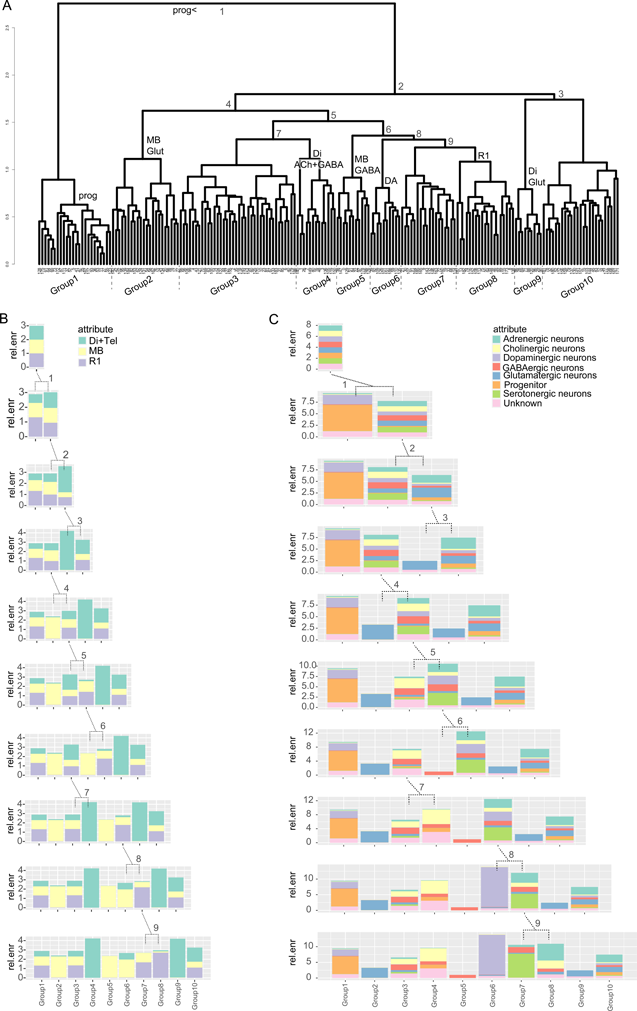
Analysis of the hierarchical tree of cell clusters. A. Hierarchical tree of scATAC-seq clusters (also shown in Figure 5). The first 10 branching levels are indicated in numbers at the point of new branch appearing. The brain region or NT-type label is shown for the cluster groups of uniform brain region or NT-type identity, at the first appearance of the cluster group. B. Relative enrichment (rel.enr) of cells originating from the different brain region samples at each indicated branch of the clusters tree. Relative enrichment is shown for the first 10 branching levels. rel.enr = 1 equals no enrichment of attribute between the groups in the given branching level, and any deviation from 1 shows either increase or decrease in relative proportion of the attribute (See Methods). The cell groups at the the brancing level 10 (Group 1 - Group 10) are shown in the tree in (A) and below the last plots in (B) and (C). C. Relative enrichment of NT-type categories in the cell groups at the first 10 branching levels. NT, neurotransmitter type; reg, brain region identity. Brain regions: Di+Tel, diencephalon and telencephalon samples; MB, midbrain samples; R1, rhombomere 1 samples. ACh+GABA, GABA- and acetylcholinergic dual neurotransmitter neurons; GLUT, glutamatergic neurons; GABA, GABAergic neurons; DA, dopaminergic neurons; prog, progenitor cells.

**Supplementary Figure 6.**
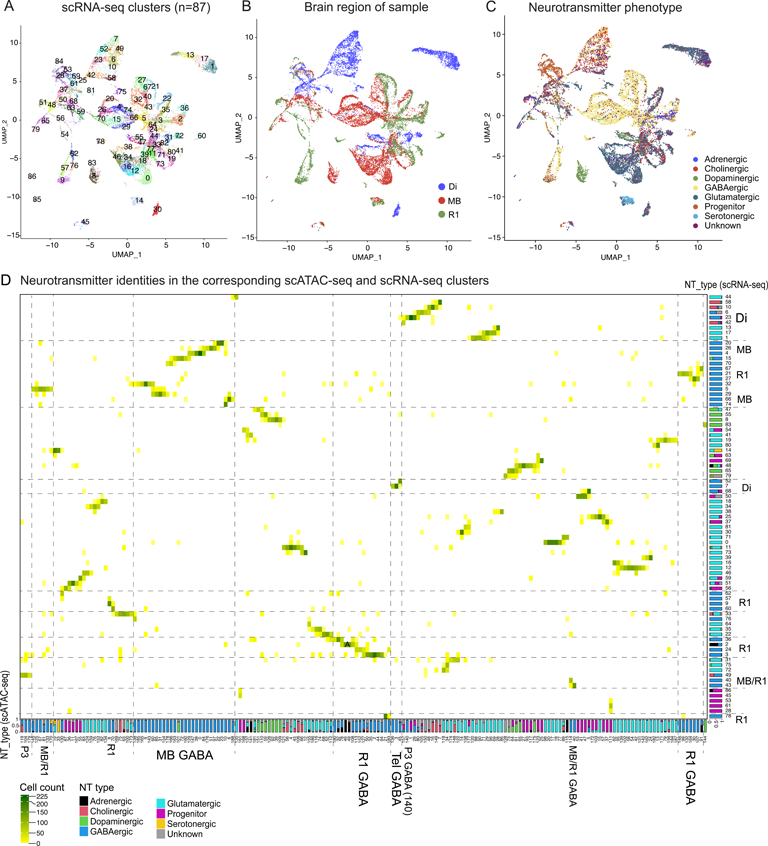
Clustering and neurotransmitter phenotype annotation of E14.5 DI, MB and R1 scRNA-seq data. A. UMAP of the 87 scRNA-seq clusters, at the optimal clustering resolution according to chooseR. B. The scRNA-seq cluster UMAP, with brain region shown for each cell. C. The scRNA-seq cluster UMAP, with neurotransmitter phenotype (NT-type) labels shown. D. Heatmap of cluster-to-cluster matches between the integrated and scATAC-seq based (columns) and scRNA-seq based (rows) clusters. The stacked barplots show the proportion of cells of each neurotansmitter type in the cluster. Matching clusters mostly received the same NT-type label and originate from the same brain region. Matching adrenergic neuron (A) clusters 2 (scRNA) vs 46, 49 (scATAC) are grouped with R1 GABAergic neurons.

**Supplementary Figure 7.**
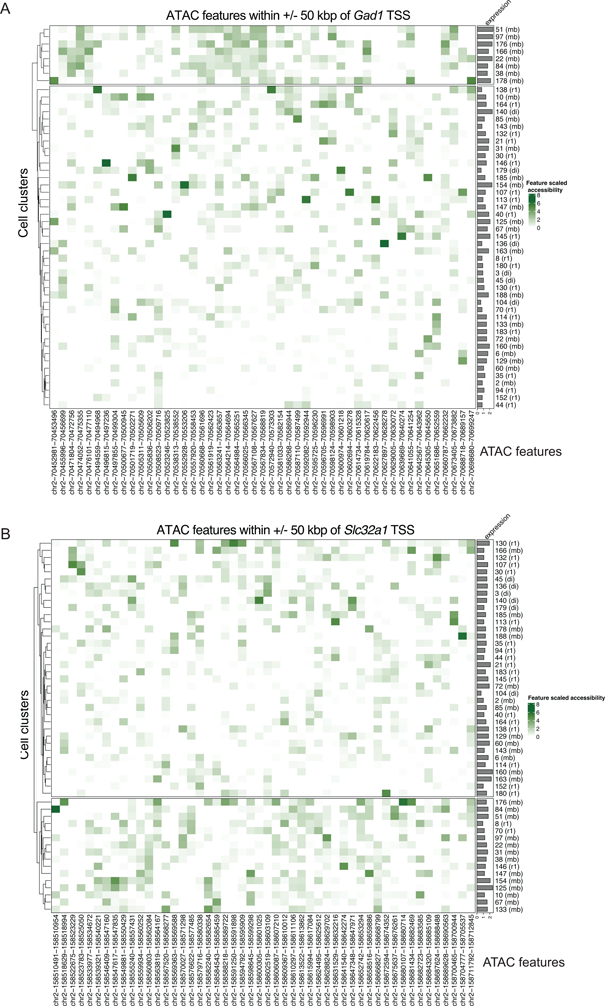
Chromatin feature accessibility around Gad1 and Slc32a1 genes in the GABAergic clusters. A Heatmap of the accessibility of ATAC features +/-50kpb around *Gad1* gene, one of the strongest indicator genes for GABAergic fate, across GABAergic clusters (rows). Dominant brain region of origin is shown in parenthesis after the cluster number. Barplots on the right side show average expression of *Gad1* in the clusters. Accessibility data has been column scaled and rows are clustered with Euclidean as distance and ward.D2 as linkage method. B. Accessibility of ATAC features within +/-50kpb of *Slc32a1* gene and the *Slc32a1* RNA expression in the GABAergic cell clusters, similar to (A).

## Supplementary Table legends

**Supplementary Table 1. Genomic position of ATAC features.**

Comparison of the position of the E14.5 ATAC features defined herein with randomised genomic position.

**Supplementary Table 2. Single-cell sample statistics.**

Single-cell RNA-seq and single-cell ATAC-seq sample statistics. The method is abbreviated in the table as ATAC, scATAC-seq or RNA, scRNA-seq. For scATAC-seq samples, the embryonic stage (E14.5), brain region of sample, genotype of the embryo, number of cells per sample, median number of fragments per cell, minimum number of fragments per barcode required to assign the barcode to a cell, the percent of reads from the sample included in final merged data, the sample name used in the SRA repository, the sample collection date (batch) and the median of ratios of fragments per cell included in the in E14.5 features is shown. For scRNA-seq samples, the embryonic stage (E14.5), brain region of sample, genotype of the embryo, number of cells per sample, median number of reads per cell, median number of genes detected in the cells, the percent of reads from the sample included in final merged data, the sample name used in the SRA repository, the sample collection date (batch) is shown.

**Supplementary Table 3. Neurotransmitter identity phenotyping.**

Sheet 1: ScType database for assigning the NT type labels.

Sheet 2: CellCycleScore genes expressed in the E14.5 mouse neuronal progenitors and list of genes used to define the *g2m* and *s* artificial genes in ScType.

**Supplementary Table 4. Cell counts per brain region and neurotransmitter phenotype.**

Identities were assigned per cell. The cluster was counted in category when >50% of cells in the cluster match the category.

**Supplementary Table 5. Unique marker gene combinations for the scATAC-seq clusters.**

Table listing the scATAC-seq cell clusters (n=189), the unique marker gene combination for each cluster, calculated using CombiROC (marker.combination), the main neurotransmitter type in the cluster (top_nt_type) and main brain region identity in the cluster (top_brain_region). The top_nt_type and top_brain_region label was assigned if >50% of the cells in the cluster receive the label, otherwise NA. The clusters labelled NA may to contain several cell types, or migratory cells found in neighboring brain regions.

**Supplementary Table 6. Clustering resolution optimization.**

Results of *ChooseR* iteration for finding optimal resolution of clustering the integrated single-cell dataset based on accessible chromatin features. The clustering was iterated from res=1 to res=20.

**Supplementary Table 7. Biological function of genes associated with differentially accessible chromatin between neighboring clusters in pseudotime.**

Analysis of Genome Ontology term enrichment in genes near differentially accessible chromatin features between cluster pairs. Selected clusters represent lineage related telencephalic and diencephalic GABAergic neurons.

**Supplementary Table 8. Comparison of scATAC and scRNA clusters.**

Contingency table of cell placement in scATAC clusters and scRNA clusters.

**Additional File at Github. Analysis of the *CombiROC* cluster marker gene expressions.** Available at https://github.com/ComputationalNeurogenetics/NeuronalFeatureSpace/blob/main/analysis/Combiroc_avg_plots110923_comp.pdf

## Notes

https://github.com/ComputationalNeurogenetics/NeuronalFeatureSpace

### Competing Interest Statement

The authors have declared no competing interest.

### Summary of Updates

The manuscript is currently under peer review and has been revised. Results of the batch effect analysis for single-cell sequencing samples have been added and the sample selection strategy and clustering quality improved.

